# Gradients of functional connectivity in the mouse cortex reflect neocortical evolution

**DOI:** 10.1101/2020.03.04.976860

**Authors:** Julia M. Huntenburg, Ling Yun Yeow, Francesca Mandino, Joanes Grandjean

## Abstract

Understanding cortical organization is a fundamental goal of neuroscience that requires comparisons across species and modalities. Large-scale connectivity gradients have recently been introduced as a data-driven representation of the intrinsic organization of the cortex. We studied resting-state functional connectivity gradients in the mouse cortex and found robust spatial patterns across four data sets. The principal gradient of functional connectivity shows a striking overlap with an axis of neocortical evolution from two primordial origins. Additional gradients reflect sensory specialization and aspects of a sensory-to-transmodal hierarchy, and are associated with transcriptomic features. While some of these gradients strongly resemble observations in the human cortex, the overall pattern in the mouse cortex emphasizes the specialization of sensory areas over a global functional hierarchy.

**Highlights:**

- The principal gradient of functional connectivity in the mouse cortex recapitulates an axis of neocortical evolution from archicortex and paleocortex.
- Additional gradients highlight sensory specialization and reflect aspects of a sensory-to-transmodal hierarchy.
- Functional connectivity gradients partly align with gene expression patterns.
- Mouse cortical gradients are stable across data sets.

## Introduction

A crucial task in neuroscience is to compare and connect findings across species. This approach enables us to learn how different cortices – including our own – have evolved and acquired their unique capacities. A major challenge for the translation of findings from animal studies to the human brain is that most techniques used in neuroscience research are too invasive to be applied in humans. A notable exception is magnetic resonance imaging (MRI), which can safely be used to investigate human brain structure and function, and is increasingly applied in small animals. MRI has the potential to bridge between studies in humans and those in established experimental species, where complementary, highly specific measures are available.

Large-scale cortical gradients were recently put forward as an analytic tool to capture the intrinsic dimensions of cortical organization (Haak et al., 2017; Huntenburg et al., 2018; Margulies et al., 2016). In contrast to parcellation approaches, which emphasize the discreteness of cortical regions or networks (Glasser et al., 2016; Yeo et al., 2011), cortical gradients focus on the significance of overarching spatial patterns in cortical features. The gradient approach was initially applied to functional connectivity data derived from resting state functional magnetic resonance imaging (rsfMRI) in the human cortex. Human functional connectivity gradients have been shown to reflect functional specialization (Haak et al., 2017; Margulies et al., 2016; Peer et al., 2019; Wang et al., 2020), align with proxies of microstructural differentiation (Huntenburg et al., 2017; Larivière et al., 2019) and predict behavioral symptoms in individuals with autism spectrum disorder (Hong et al., 2019).

Beyond being simply another approach to analyze rsfMRI data, the relevance of cortical gradients derives from their roots in classic theories of cortical organization and evolution. Early neuroanatomical studies described gradients in cortical microstructure and connectivity across species, and postulated that they represent the course of cortical evolution.Specifically, the dual origin theory states that the neocortex (or isocortex) evolved in successive waves emanating from two allocortical origins: the archicortex (hippocampus) and the paleocortex (piriform cortex) (Abbie, 1942, 1940; Dart, 1934; Sanides, 1970, 1962). The dual origin theory entails a spatial organization in which evolutionary newer areas emerge further and further away from the two origins, forming a spatial gradient of neocortical evolution. This organization is reflected in gradients of progressive microstructural differentiation, with more recently evolved areas showing less laminar differentiation.Connectivity patterns have been found to align with these microstructural gradients, as cortico-cortical connections preferentially occur between areas that occupy similar gradient positions (Barbas and Pandya, 1989; Pandya et al., 2015; Pandya and Sanides, 1973; Pandya and Yeterian, 1985). Based on this legacy, one core aim of contemporary work on cortical gradients is to use modern methods to describe and compare gradients across species. Which fundamental aspects of cortical organization can be found across cortices that widely vary in size and shape? To what extent do gradients represent species-specific organization?

Recent studies in non-human species have described cortical gradients based on tract-tracing connectivity data in the mouse, cat, marmoset and macaque monkey (Buckner and Margulies, 2019; Fulcher et al., 2019; Goulas et al., 2019b, 2019a; Margulies et al., 2016). Many of these studies included additional cortical features such as cytoarchitectonic differentiation or gene expression patterns. Their findings are compelling as they illustrate both similarities and differences across species, and highlight the potential cellular and molecular basis of cortical gradients. One potential limitation of this line of work is that it relies on post-mortem data, likely making it less sensitive to organizational features that reflect different functional modes. In addition, the data is typically averaged across predefined cortical parcels which prevents the detection of spatial patterns that do not adhere to predefined areal boundaries.

In the present study, we complement previous work on gradients in the mouse cortex with an analysis of functional connectivity gradients in vivo. In contrast to most human studies, high-quality rsfMRI in mice requires acquisition in a lightly anesthetized state. Here, we use four mouse rsfMRI data sets, acquired under an optimized anesthesia protocol, and present robust cortical gradients. In an exploratory approach, we inspect these functional connectivity gradients with respect to their potential functional significance and their relationship to previously described gradients in structural features.

## Material and methods

### Data sets

Throughout this study we used the C57Bl6 control animals of four publicly available data sets (*see Data and code availability*). The primary analyses, displayed in Figures 1–4, were performed on data from 10 female mice (age = 3 months), referred to as *Main* where necessary. For the robustness analyses we used three additional data sets, referred to as *AD2* (Grandjean et al., 2016b), *AD3* (Mandino et al., 2019) and *CSD1* (Grandjean et al., 2016a), after the naming convention of the underlying database. We used data from 18 female and 14 male mice (age = 13 months) from AD2, 10 male mice from AD3 (age = 3/6 months in session 1/2) and 51 male mice from CSD1 (age = 3 months).

**Fig. 1.**
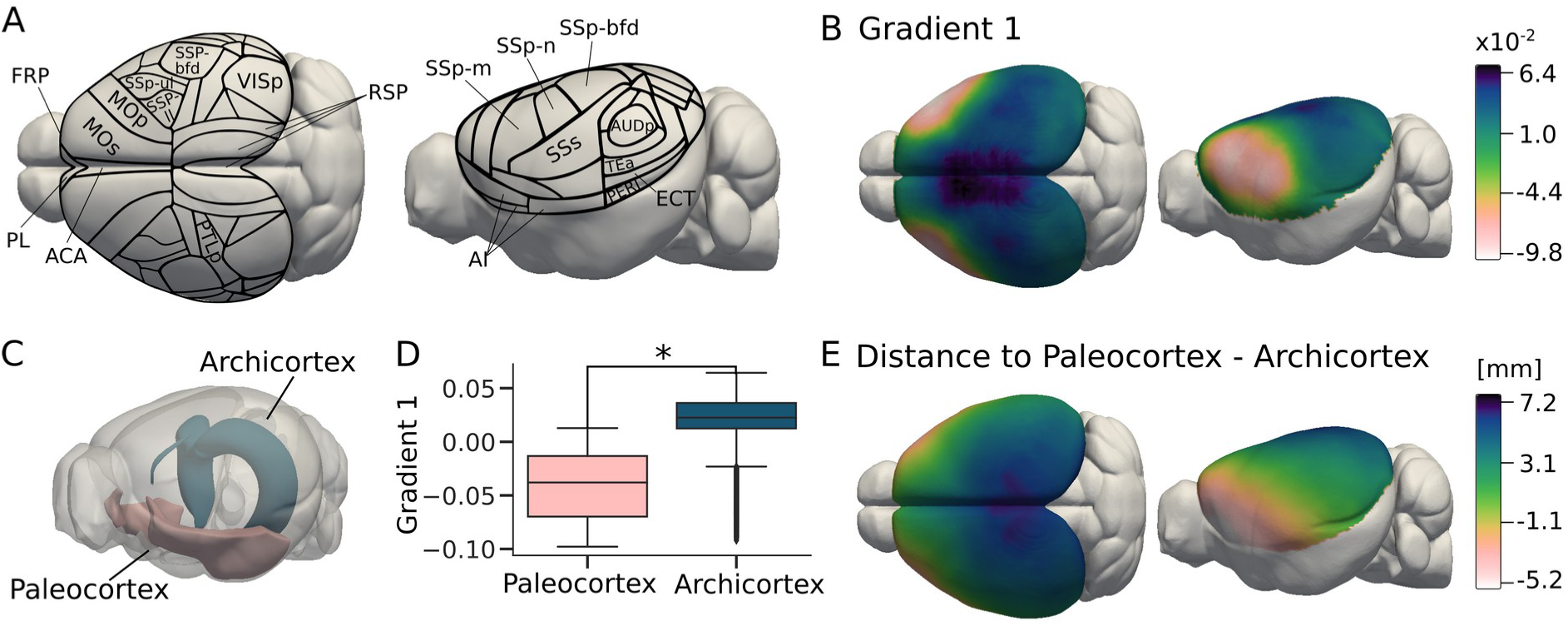
The principal gradient and the two neocortical origins. A) Cortical parcellation based on the Allen Brain Mouse Atlas. B) The principal gradient of functional connectivity sampled on the surface mesh. Note that the gradients are abstract dimensions and have no units. C) Mesh representations of the hippocampal area (archicortex, *petrol*) and piriform area (paleocortex, *rose*), that were used as source regions to calculate geodesic distance to the two neocortical origins. D) The cortical surface was divided into two zones based on the minimal distance of each surface node to either paleocortex or archicortex. Shown are the Gradient 1 values in either zone (Wilcoxon rank-sum = 69.6, P_corr_ = 0.01). E) Map of the combined geodesic distance, obtained by subtracting distance to archicortex from distance to paleocortex. High negative values indicate proximity to paleocortex, high positive values indicate proximity to archicortex. Values close to zero mark that a surface node is equally distant from both origins.

**Fig. 2.**
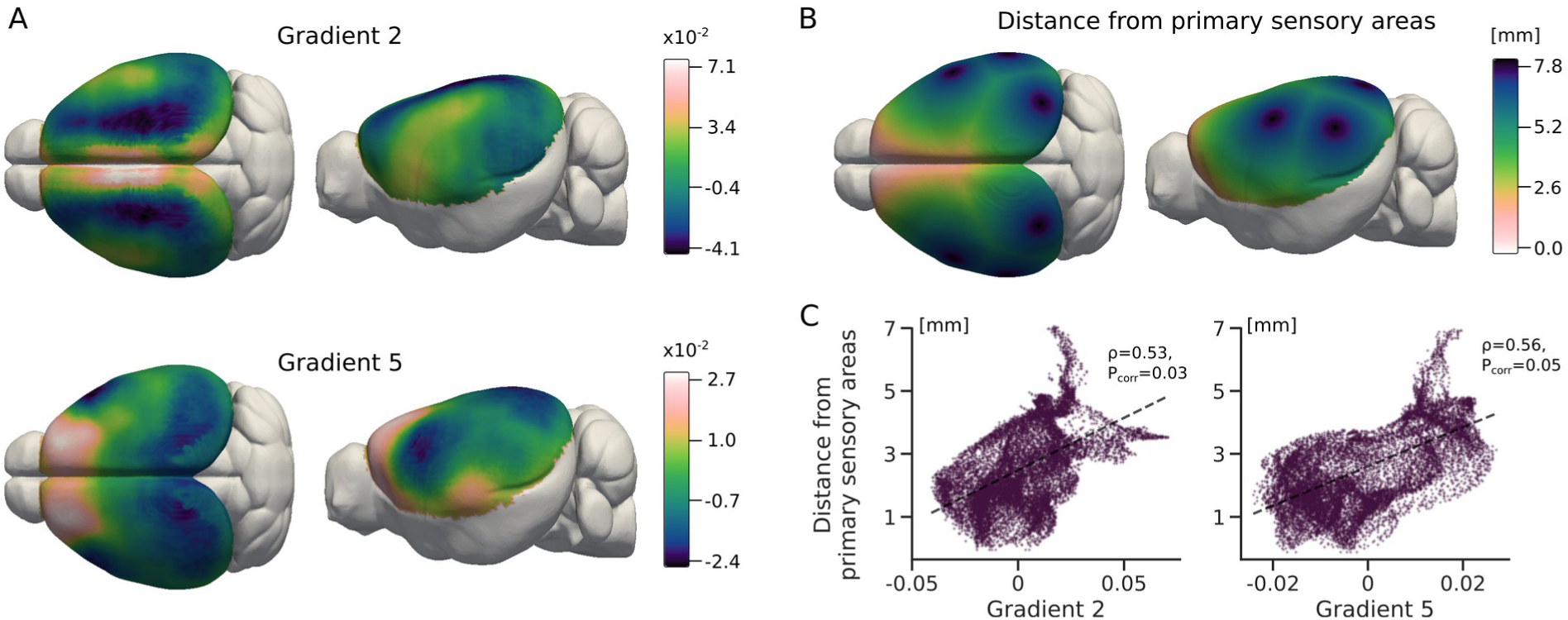
Sensory-to-transmodal gradient patterns. A) Functional connectivity Gradient 2 and 5 show a general sensory-to-transmodal organization. Geodesic distance from primary sensory centers (B) is correlated to Gradient 2 and 5 (C).

**Fig. 3.**
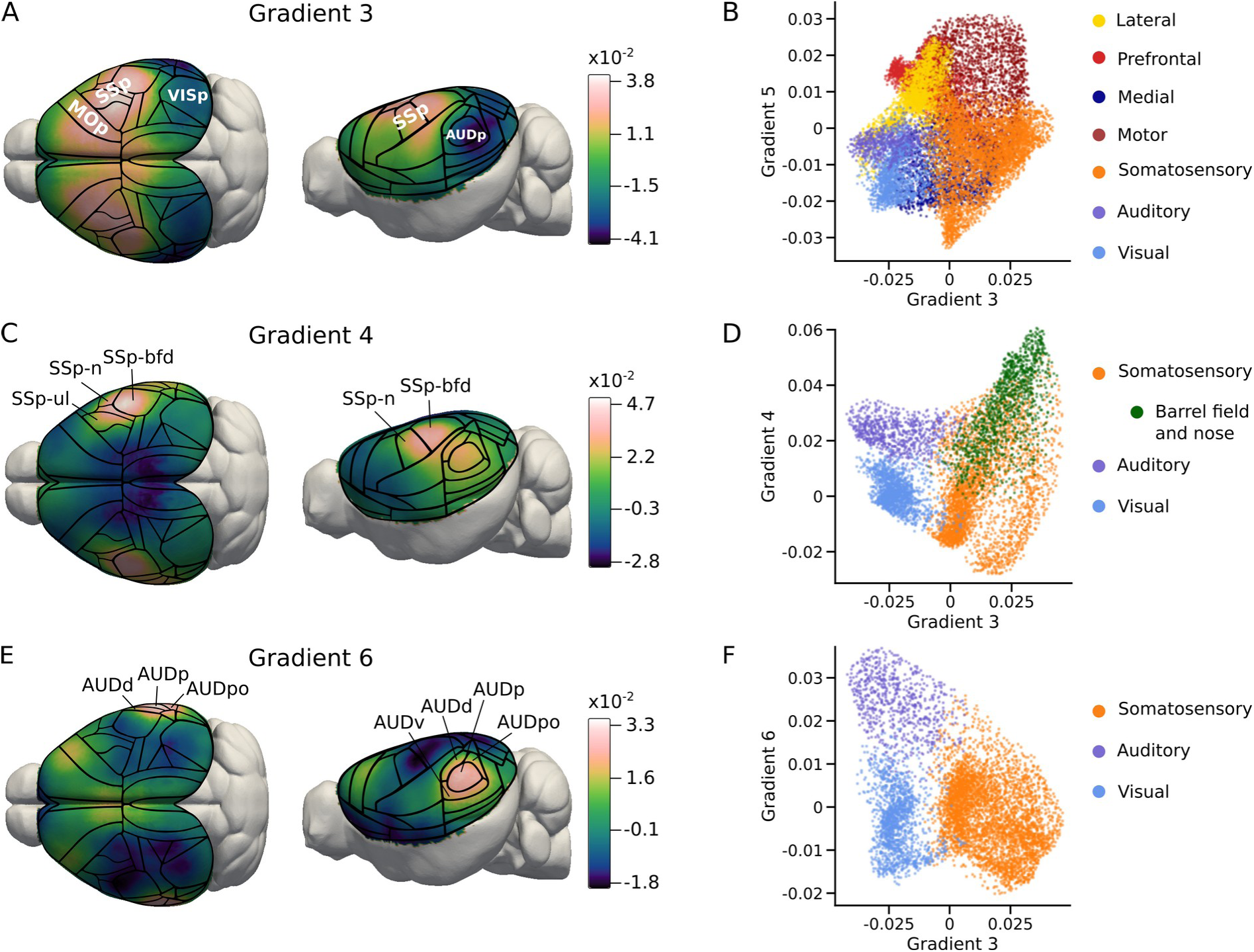
Gradients of sensory specialization. A) Gradient 3 reflects a dissociation between somatomotor cortex and auditory/visual areas. C) Gradient 4 highlights the barrel-field and nose area of the primary somatosensory cortex. E) Gradient 6 separates auditory areas from the rest of the cortex. B,D&F) Scatter plots of different gradient combinations help to visualize the aspects of functional organization captured in these gradients. The cluster assignment has been adapted from Harris et al. (2019). In order to highlight features of sensory specialization, only sensory clusters are plotted in D) and F).

**Fig. 4.**
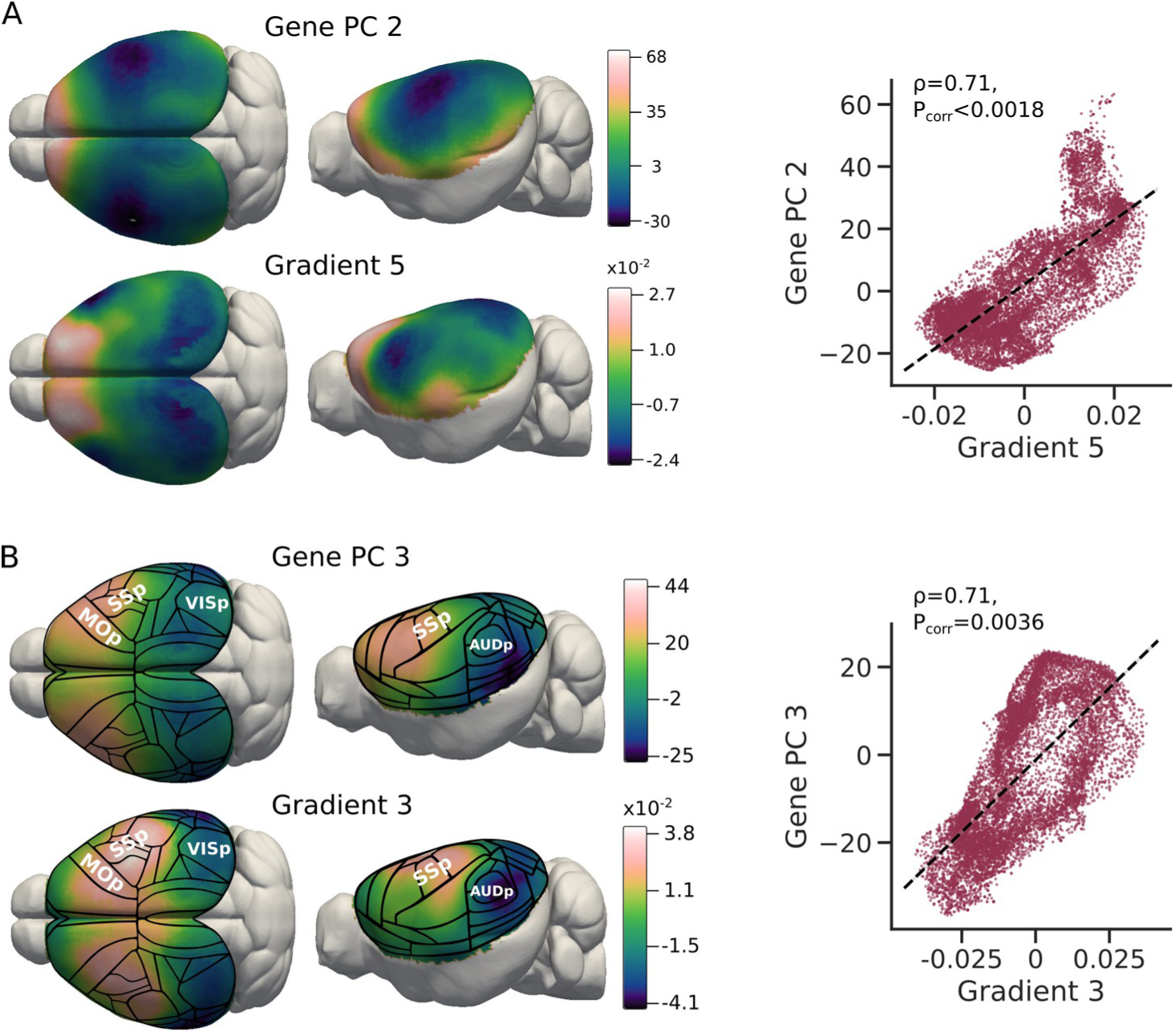
Correlation to dominant gene expression patterns. Gene PC shows a strong spatial correlation to sensory-to-transmodal Gradient 5 (A). Gene PC3 strongly resembles somatomotor-audiovisual Gradient 3 (B).

### Animal preparation for imaging

All applicable international, national, and/or institutional guidelines for the care and use of animals were followed. All procedures performed in studies involving animals were in accordance with the ethical standards of the Institutional Animal Care and Use Committee (A*STAR Biological Resource Centre, Singapore, IACUC #161134/171203). Animal preparation for all data sets was performed as described in Grandjean et al. (2014).Anesthesia was induced with 4% isoflurane; subsequently, the animals were endotracheally intubated, placed on a MRI-compatible cradle and artificially ventilated (90 breaths/minute; Kent Scientific Corporation, Torrington, Connecticut, USA). A bolus with a mixture of medetomidine (Dormitor, Elanco, Greenfield, Indiana, USA) and Pancuronium Bromide (muscle relaxant, Sigma-Aldrich Pte Ltd, Singapore) was administered subcutaneously (0.05 mg/kg), followed by a maintenance infusion (0.1 mg/kg/h) while isoflurane was simultaneously reduced and kept to 0.5%. Care was taken to maintain the animal temperature at 37°C.

### Data acquisition

Data for the Main data set were acquired on an 11.75 T (Bruker BioSpin MRI, Ettlingen, Germany) equipped with a BGA-S gradient system, a 72mm linear volume resonator coil for transmission and a cryoprobe 2×2 phased-array surface coil. Images were acquired using Paravision 6.0.1 software. An anatomical reference scan was acquired using a spin-echo Turbo-RARE sequence: field of view (FOV) = 17 × 9 mm^2^, FOV saturation slice masking non-brain regions, number of slices = 28, slice thickness = 0.35, slice gap = 0.05 mm, matrix dimension (MD) = 200 × 100, repetition time (TR) = 2742 ms, echo time (TE) = 30 ms, RARE factor = 8, number of averages = 2. fMRI was acquired using a gradient-echo echo-planar imaging (GE-EPI) sequence with the same geometry as the anatomical: MD = 90 x 60, in-plane resolution = 0.19 x 0.15 mm^2^, TR = 1000 ms, TE = 11.7 ms, flip angle = 50°,volumes = 180, bandwidth = 119047 Hz. Field inhomogeneity was corrected using MAPSHIM protocol. Ten animals were imaged over two sessions, each consisting of three consecutive GE-EPI runs, resulting in 60 runs in total. For the AD3 data set, scanner and imaging parameters were equal to *Main*, expect for a slightly longer TE (15 ms) and run length (600 volumes). All 10 animals were imaged in two sessions, acquired three months apart. Each session contained a single GE-EPI run, resulting in 20 runs in total.

Data for AD2 and CSD1 were acquired on a 9.4 T scanner (Bruker BioSpin MRI, Ettlingen, Germany) equipped BGA-S gradient system, a linear volume resonator coil for transmission, and a cryoprobe 2×2 phased-array surface coil. GE-EPI imaging parameters for both data sets were MD = 90×70, in-plane resolution = 0.22×0.25mm^2^, TR = 1000 ms, TE = 9.2 ms, flip angle = 90°, volumes = 360, bandwidth = 250000 Hz. For AD2 all 32 animals were imaged in a single session with a single GE-EPI run, resulting in 32 runs in total. For CSD1, 25 animals were imaged in one session and 26 animals were imaged in two sessions, of one run each, resulting in 77 runs in total.

### Data preprocessing

A study-specific template was created from the individual anatomical images using the *buildtemplateparallel*.*sh* script from the Advanced Normalization Tools (ANTs, 20150828 builds, Avants et al., 2011). The study template was subsequently registered to the reference template of the Allen Mouse Common Coordinate Framework version 3 (CCF v3) (Dong, 2008; Lein et al., 2007; Oh et al., 2014). Individual anatomical images were corrected for b1 field inhomogeneity (*N4BiasFieldCorrection*), denoised (*DenoiseImage*), intensity thresholded (*ImageMath*), brain masked (*antsBrainExtraction*.*sh*), and registered to the study template, resampled to a 0.2 x 0.2 x 0.2 mm^3^ resolution, using a SyN diffeomorphic image registration protocol (*antsRegistration*). The functional EPI time series were despiked (*3dDespike*) and motion-corrected (*3dvolreg*) using the Analysis of Functional NeuroImages software (AFNI_16.1.15, Cox, 1996). A temporal average was estimated using *fslmaths* (FMRIB Software Library 5.0.9, Jenkinson et al., 2012), corrected for the b1 field (*N4BiasFieldCorrection*), brain masked (*antsBrainExtraction*.*sh*), and registered to the anatomical image (*antsRegistration*). The fMRI times series were bandpass filtered (*3dBandpass*) to a 0.01 to 0.25 Hz band. An independent component analysis (*melodic*) was performed and used as a basis to train a *FIX* classifier (Salimi-Khorshidi et al., 2014) in 15 randomly selected runs. The classifier was applied to identify nuisance-associated components that were regressed from the fMRI time series. The EPI to anatomical,anatomical to study template, and study template to Allen Mouse CCF v3 registrations were combined into one set of forward and backward transforms (*ComposeMultiTransform*). The combined transforms were applied to the denoised fMRI data (Fig. S1).

### Dimensionality reduction of rsfMRI data

Resting state time series of each rsfMRI run were masked to the isocortex using a downsampled mask from the Allen Mouse CCF v3. Data were smoothed within that mask with a Gaussian kernel (FWHM = 0.45 mm) using Nilearn (Abraham et al., 2014). A functional connectivity matrix was derived by computing the Pearson product-moment correlation between the time series of each pair of isocortical voxels. The individual matrices from all runs were Fisher r-to-z-transformed, averaged across all runs, and back-transformed to Pearson’s r values. The average functional connectivity matrix was decomposed through nonlinear dimensionality reduction using diffusion maps (Coifman and Lafon, 2006). Diffusion maps have been shown to be well-suited for the analysis of human resting-state fMRI data,e.g. in the context inter-subject alignment (Langs et al., 2015b, 2015a, 2014, 2010) or to study global cortical organization (Hong et al., 2019; Huntenburg et al., 2017; Larivière et al., 2019; Margulies et al., 2016). In contrast to linear methods, nonlinear decomposition approaches can retain both local and long-range connectivity relationships in the embedded space, without requiring kernel manipulations. Diffusion maps in particular are robust to noise in the similarity matrix, and stable with regards to its size (Lafon and Lee, 2006).

In brief, the average functional connectivity matrix was transformed into a positive similarity matrix ***L*** by increasing each element by 1 and dividing it by 2.The similarity matrix was then transformed into a Markov chain with transition probability matrix **M** as follows:

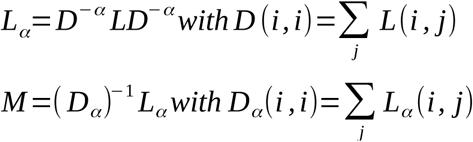

The diffusion operator α can range between 0 and 1, in the latter case **M** is equal to the normalized graph Laplacian. In keeping with previous rsfMRI studies, we chose a diffusion operator of α=0.5. Thereby, transition in the Markov chain approximates a Fokker–Planck diffusion process, which renders the method less sensitive to nonuniform sampling of the data on the underlying manifold (cf. Coifman and Lafon, 2006, for an in-depth discussion). The eigenvectors of **M** represent the coordinate system of the embedded space. Here we embed the 15k x 15k matrix in 100 dimensions, to which we refer as functional connectivity gradients. Each one-dimensional gradient captures a distinct aspect of functional connectivity patterns in the original matrix. The gradients are ordered by decreasing eigenvalue, i.e. they capture a decreasing amount of variance in the Markov transition matrix. When plotting the variance captured by each individual gradient, the first clear “elbow” is visible at Gradient 6 (Fig. S2). We therefore focus on Gradient 1-6 for the remainder of this study. These gradients were projected back onto the isocortical voxels and sampled on a surface mesh representation of the Allen Mouse CCF v3 for visualization. For comparison, we also carried out a linear decomposition of the average functional connectivity matrix in 100 dimensions, using principal component analysis (PCA) as implemented in scikit-learn (Pedregosa et al., 2011).

### Surrogate maps and statistical testing

Like most brain maps, functional connectivity gradients show a substantial degree of spatial autocorrelation (SA), violating the assumption of independence in standard statistical tests. We addressed this issue by implementing permutation tests based on SA-matched surrogate maps. The surrogate maps were created using a generative model recently put forward by Burt et al. (2020). This approach is based on matching SA-variograms (i.e. the variance between pairs of data points as a function of their distance) between surrogate and target maps, and provides support for volumetric and surface maps. We generated 10,000 surrogate maps for each Gradient 1-6, in both volumetric and surface space. All statistical analyses were performed by calculating the respective test statistic for a gradient and its surrogates, and calculating the p-value as the incidence of test statistics equal to, or more extreme than, that obtained for the gradient in question. Where applicable, we used Bonferroni correction for multiple comparisons, e.g. when testing across all six gradients.Unless explicitly stated, all P-values are reported after Bonferroni correction (indicated P_corr_).

### Geodesic distance from origins and primary sensory regions

In order to investigate the spatial layout of the functional connectivity gradients, we operationalized spatial relationships to areas of interest as geodesic distance along the cortical surface. Geodesic distance measures the length of the shortest line between two points on a surface, such that the line lies on the surface. We calculated the geodesic distance based on the surface mesh representation of the Allen Mouse CCF v3 using the gdist package provided by the Virtual Brain Project (Leon et al., 2013).

For the dual origin analysis, we used volumetric masks of the piriform area and the hippocampal region from the Allen Mouse CCF v3 (Fig. 1C), and sampled these masks onto the surface mesh. We then created two distance maps by assigning to each isocortical surface node its shortest distance to any node within the hippocampal and piriform mask, respectively. We divided the cortex into two zones, where each surface node was assigned to an “archicortical” or “paleocortical” zone, depending on whether it had a shorter distance to the hippocampus or piriform area. To assess whether Gradient 1-6 values in the two zones were likely to be drawn from different distributions, we computed the Wilcoxon rank-sum statistic as implemented in Scipy (Virtanen et al., 2020). We also created a combined distance map by subtracting the hippocampal distance map from the piriform distance map (Fig. 1E). The combined distance map captures a spatial gradient between the two origins at the extreme ends of the value scale. Concretely, high positive values indicate proximity to hippocampus and high negative values indicate proximity to the piriform area (the directionality is arbitrarily chosen and could also be flipped). Values close to zero indicate maximum distance from both origins. The spatial correlation between this combined distance map and Gradient 1-6 was computed as Spearman’s rank correlation. Statistical significance for the rank-sum test and the spatial correlation was assessed using permutation tests as described above. P-values were adjusted for multiple comparison across the six gradients using Bonferroni correction.

To compute the geodesic distance to primary sensory regions, we projected masks of the primary visual, auditory and somatosensory areas onto the surface mesh. We then found the surface node closest to the geometric center of each primary sensory region for each hemisphere, resulting in six sensory center nodes in total. We calculated a distance map in which each isocortical surface node is assigned its shortest possible distance to any of these six sensory center nodes. The spatial correlation between this distance map and Gradient 1-6 was calculated using Spearman’s rank correlation. Statistical significance was assessed using permutation tests as described above. P-values were adjusted to account for the six simultaneous tests using Bonferroni correction.

### Gene expression

In order to extract dominant patterns of gene expression in the mouse cortex, we used 3D reconstructed data from RNA in-situ hybridization experiments, provided as part of the Allen Mouse Brain Atlas (Lein et al., 2007). Concretely, we based our analysis on the subset of 4,376 coronal experiments underlying the Anatomic Gene Expression Atlas (AGEA) (cf. Supplementary Table 5 in Ng et al., 2009).For 3,964 of these experiments, gene expression energy maps were available for download and an Entrez Gene ID was provided. These 3,964 expression energy maps were used for subsequent analyses. Since many of the energy maps do not cover the entire isocortex, we created a mask containing only isocortical voxels with missing data in less than half of all maps, and only considered data withing that mask. We z-scored the expression energy maps within this mask, averaged maps of experiments probing the same gene (as per Entrez Gene ID), and normalized the data using a sigmoid transfer function (Fulcher and Fornito, 2016), and standard scaling. A PCA was performed on the standardized data. Based on the elbow in the variance distribution we kept the first four principal components (Gene PC1-4) and projected them back into the voxel space. Gene PC1 showed a clear cortical depth-dependent structure, indicating that it captures layer-specific gene expression (Fig. S3). Our rsfMRI data is not well-suited to reveal such structure in the functional connectivity data, due to the spatial resolution of the BOLD response, its bias to superficial layers and the smoothing we applied to the data. We therefore disregarded Gene PC1 from the following analysis. Gene PC2-4 were projected onto the surface and correlated with the six functional connectivity gradients using Spearman’s rank correlation. Statistical significance was assessed through permutation testing as described above. P-values were adjusted to account for 18 pairwise correlations using Bonferroni correction. Gene ontology (GO) enrichment analysis was performed by ranking all genes according to their loading onto Gene PC2 and 3, and using these ranked lists as input for the GOrilla tool (Eden et al., 2009). Enriched GO terms that appeared densely at the top of the ranked list were identified by GOrilla’s algorithm. Correction for multiple comparison across 11,266 GO terms in the database was performed using the Benjamini and Hochberg (1995) method, controlling the false discovery rate (FDR) at 5%.

### Cytoarchitectonic differentiation

To assess the relationship of functional connectivity gradients to cytoarchitectonic differentiation we adopted an ordinal scale of cortical types from Goulas et al. (2016). This scale is based on expert annotation of isocortical areas according to their cytoarchitectonic features in high-resolution Nissl stained sections (Paxinos and Franklin, 2013; Van De Werd and Uylings, 2014). The ordinal scale of cortical type ranges from 1 in agranular areas with a rather undifferentiated laminar pattern, to 4 in highly differentiated, eulaminate areas with a pronounced layer IV (see Goulas et al., 2016 for more details). In order to relate this measure to the functional connectivity gradients, we averaged the values for each gradient within each cortical area (derived from the Allen Mouse CCF v3) and computed Kendall’s tau-b rank correlation for ordinal measures between gradient values and cortical type. Statistical significance was assessed using permutation tests as described above. P-values were adjusted for the simultaneous tests across six gradients using Bonferroni correction.

### T1w:T2w MRI

The ratio of T1-weighted (T1w) over T2-weighted (T2w) MR images has been put forward as a readily available proxy for the spatial distribution of intracortical myelin (Glasser and Van Essen, 2011). In order to probe the relationship between the functional connectivity gradients and this measure, we obtained the post-mortem T1w and T2w MRI images that are distributed as part of the Waxholm space (Johnson et al., 2010). The T1w and T2w images were corrected for b1 field inhomogeneity (*N4BiasFieldCorrection*), registered to the Allen Mouse CCF v3 template *(antsRegistration)*, downsampled to match the resolution of the functional connectivity gradients and masked to the isocortex. We computed the ratio of T1w over T2w values within this mask and computed Spearman’s rank correlation of the T1w:T2w map to Gradient 1-6. Statistical significance was assessed using permutation testing as described above and P-values were adjusted for simultaneous testing across the six gradients using Bonferroni correction.

### Data and code availability

All in vivo MRI data used in this study are available through public data repositories, in raw and preprocessed form (Table S1). The Allen Mouse CCF v3 template, surface mesh and structure masks, as well as the Allen Mouse Brain Atlas gene expression maps were obtained through the Allen Software Development Kit (Allen SDK, https://github.com/AllenInstitute/AllenSDK). The Waxholm space T1w and T2w data were downloaded from the Scalable Brain Atlas (https://scalablebrainatlas.incf.org/mouse/WHS12). All code used for the preparation of this manuscript is available on Github at https://github.com/grandjeanlab/MouseMRIPrep (preprocessing) and https://github.com/juhuntenburg/mouse_gradients (analysis and visualization).

## Results

To investigate the intrinsic functional organization of the mouse cortex, high quality rsfMRI data were obtained in ventilated, lightly-anesthetized mice. After careful preprocessing (Fig. S1), a functional connectivity matrix was calculated and decomposed into a set of one-dimensional gradients, each capturing part of the variance in functional connectivity patterns across the cortex. Each gradient can be thought of as a spectrum of functional connectivity similarity: Cortical locations that have similar Gradient *x* values resemble each other in the aspect of functional connectivity that is captured in Gradient *x*. Cortical locations positioned on opposite extremes of Gradient *x* differ maximally in this aspect of functional connectivity. The curve of the variance captured by individual gradients flattens after Gradient 6 (Fig. S2). We therefore decided to focus our analyses on Gradient 1-6.

### The principal gradient of functional connectivity reflects spatial distance from neocortical origins

The principal gradient – or Gradient 1 – captures the highest amount of variance in the functional connectivity data (Fig. S2). Upon visual examination, we noticed that its spatial distribution showed are marked similarity to previous representations of the dual origin organization in the mouse cortex (Fig. 1B, cf. Goulas et al., 2019b). We therefore tested the hypothesis that the principal gradient reflects a spatial progression from the two postulated neocortical origins, the archicortex (hippocampus) and paleocortex (piriform area) (Fig. 1C). To this end, we computed the geodesic distance along the cortical surface from the hippocampus and piriform cortex. First, we divided the cortex into a paleocortical and an archicortical zone, based on the minimal distance of every surface node to either origin. We found that Gradient 1 values differ significantly between the two zones (Wilcoxon rank-sum = 69.6, P_corr_ = 0.01, Fig. 1D). We then created a combined distance map by subtracting distance to archicortex from distance to paleocortex (Fig 1E). In this combined distance map, values close to zero indicate maximum distance to either origin, i.e. the transition between the two zones from the previous analysis. Large negative values indicate proximity to archicortex, large positive values indicate proximity to paleocortex. Gradient 1 shows a strong spatial correlation to this combined distance from the two cortical origins (ρ = 0.82, P_corr_ = 0.01). We repeated the analyses for the remaining five gradients but found neither a significant difference between the two zones, nor a significant spatial correlation to the combined distance maps for Gradients 2-6. These results demonstrate a close link between the principal gradient of functional connectivity and distance from the two origins of the neocortex.

### Gradients 2 and 5 are associated with distance from primary sensory regions

Studies of the human cortex have consistently found the principal gradient of functional connectivity to span between unimodal sensorimotor areas at one end and transmodal association regions at the other (Hong et al., 2019; Huntenburg et al., 2017; Margulies et al., 2016). We explored the current data set for similar global patterns in the mouse cortex and found two functional connectivity gradients, Gradient 2 and 5, that capture two aspects of a sensory-to-transmodal cortical progression (Fig. 2A). The transmodal extreme end of Gradient 2 lies in medial and prefrontal regions such as the anterior cingulate cortex, retrosplenial, prelimbic and infralimbic areas, and the frontal pole. The sensory extreme on the other end comprises trunk and lower limb areas of primary somatosensory cortex as well as primary visual, auditory and motor cortex. Lateral regions such as the agranular insula, ecto- and perirhinal area fall in between these extremes. Of note, parts of the mouth, nose and barrel field primary somatosensory areas also cluster in the middle or even towards the “transmodal” end of Gradient 2, somewhat defying the simple sensory-to-transmodal description. Along Gradient 5, the sensory regions cluster more clearly towards one end of the gradient. The transmodal extreme here includes prefrontal and lateral regions, while the medial retrosplenial areas are shifted more towards the sensory regions. Another clear difference to Gradient 2 is that the motor regions fall on the “transmodal” end of Gradient 5.

In humans, the sensory-to-transmodal gradient has been shown to largely recapitulate a map of geodesic distance between primary sensory and transmodal regions (Huntenburg et al., 2018; Margulies et al., 2016). In order to test whether a similar link can be found in the mouse cortex, we created a map of geodesic distance to the geometric centers of the primary somatosensory, visual and auditory areas (Fig. 2B). Note that each surface node is assigned its *shortest possible* geodesic distance to any of the primary sensory centers. This distance map showed a significant spatial correlation with Gradient 2 (ρ = 0.53, P_corr_ = 0.03) and Gradient 5 (ρ = 0.56, P_corr_ = 0.05, Fig. 2C), but not with Gradient 1,3,4 or 6.

In summary, Gradient 2 and 5 reflect two aspects of a sensory-to-transmodal organization, with their transmodal extremes in medial (Gradient 2) and lateral (Gradient 5) regions, respectively. The spatial organization of both gradients substantially overlaps with the distance from primary sensory regions.

### Gradients 3,4 and 6 capture the sensory specialization of the mouse cortex

We observed that the remaining Gradients 3, 4 and 6 reflect a high level of specialization within sensory cortex. Gradient 3 separates somatosensory (and motor) areas at one extreme end from visual and auditory regions at the other (Fig. 3A). Together, Gradient 3 and the sensory-to-transmodal Gradient 5 build a two-dimensional coordinate system of functional organization in the mouse cortex, which captures previously established cortical clusters without imposing discrete boundaries (Fig. 3B)(clusters adapted from Harris et al., 2019). Gradient 4 and 6 capture variance associated with an even higher level of sensory specialization and emphasize some of the most important sensory inputs for the mouse. Within primary somatosensory cortex, Gradient 4 separates the barrel-field and nose areas from the trunk and lower limb areas (Fig. 3C & D). Gradient 6 singles out the auditory cortex versus all other sensory regions (Fig. 3E & F). Together, Gradient 3, 4 and 6 capture the dissociation of different sensory dimensions in the mouse cortex.

### Correlation to dominant patterns of gene expression

Our next question was if functional connectivity gradients are associated with dominant patterns of gene expression in the mouse cortex. We therefore used data from the Allen Mouse Brain Atlas (Lein et al., 2007) to extract principal components of gene expression (Gene PCs). We observed that the variance captured by each principal component flattened after Gene PC4, and excluded all further components from the analysis. Gene PC1 evidently reflected layer-specific gene expression patterns (Fig. S3). Since the rsfMRI data based functional connectivity gradients themselves do not resolve depth-dependent structure, we disregarded Gene PC1 in the comparison to the gradients. Pairwise spatial correlation of Gene PC2-4 with Gradient 1-6 revealed two significant relationships: Gene PC2 showed a substantial correlation with the sensory-to-transmodal Gradient 5 (ρ = 0.71, P_corr_ < 0.0018, Fig. 4A). Similarly, Gene PC3 was highly correlated with the somatomotor-to-audiovisual Gradient 3 (ρ = 0.71, P_corr_ = 0.0036 Fig. 4b). In an effort to gain insight into the functional relevance of Gene PC2 and 3, we performed a gene ontology enrichment analysis, based on the loading of all genes onto these two components. Only PC3 showed a significant enrichment, for gene ontologies associated with the regulation of the cell cycle, of developmental processes, of cell differentiation, and of cellular component movement (Table S2). Taken together, two dominant spatial patterns of gene expression in the mouse cortex are strongly correlated with two of the functional connectivity gradients examined in this study.

### Relationship to measures of microstructural differentiation

We also explored the relationship of Gradient 1-6 to microstructural differentiation across the mouse cortex using two metrics. First, we correlated the average gradient values of all isocortical regions with cytoarchitectonic type, assigned on an ordinal scale by Goulas et al. (2016) (Fig. S4A). We found a moderate association between cytoarchitectonic type and Gradient 2, however, this results did not remain statistically significant after correcting for multiple comparison (Kendall’s τ = −0.39, P_uncorr_ = 0.05, P_corr_ = 0.3, Fig. S4B). Second, as a continuous measure of microstructural differentiation, we used the ratio of T1-weighted over T2-weighted post-mortem MR images (T1w:T2w), derived from the Waxholm space (Johnson et al., 2010). The T1w:T2w map showed a small, but significant association with Gradient 3 (ρ = 0.19, P_corr_ = 0.018, Fig. S4C). Thus, we could observe only a weak relationship between functional connectivity gradients and measures of microstructural differentiation in the mouse cortex.

### Robustness of functional connectivity gradients

In order to assess the robustness of Gradient 1-6, we performed the same preprocessing and dimensionality reduction in three additional rsfMRI data sets: AD2, AD3 and CSD1, comprising 32, 20 and 77 functional runs, respectively. We found that all data sets showed a clear dominance of Gradient 1 in terms of variance represented (Fig. S2). AD2 and CSD1 showed an elbow in individual variance at Gradient 6, exactly like the Main data set, while AD3 had an earlier elbow at Gradient 4 (Fig. S2). The principal gradient was strikingly robust in its spatial pattern across all four data sets (Fig. S5). In order to obtain a quantitative measure of similarity across the different data sets, we computed the pairwise spatial correlation between all six gradients of the Main data set and each of the test data sets AD2, AD3 and CSD1 (Fig. S6A-C). The correlation between Gradient 1 of the Main data set and Gradient 1 of all test data sets was high (ρ = 0.89/0.96/0.81 for AD2/AD3/CSD1, P_corr_ < 0.0036 in all cases). The results for Gradients 2-6 were more variable. However, for each gradient in the Main data set, there was at least one analog gradient with a moderate-to-high, significant (P_corr_ ≤ 0.05) spatial correlation in each test data set. The only exception was the lack of an analog to Main Gradient 2 in the AD2 data set. We also tested the robustness of the gradients to the decomposition approach, by replacing diffusion maps by a linear PCA in the Main data set. The first six components of this PCA showed a very high spatial correlation to Gradient 1-6 and even reproduced their original order (Fig. S6D).

Therefore, the general aspects of the functional connectivity gradients discussed in the previous sections could be reproduced across three additional data sets, and an alternative decomposition method. Gradient 1 in particular is highly stable throughout all analyses. While their exact order varies, the spatial patterns captured in Gradients 2-6 were also reproduced across data sets.

## Discussion

We investigated the intrinsic functional organization of the mouse cortex through resting-state functional connectivity gradients. The most robust organizational feature was found to be a prominent gradient reflecting the spatial distance from the presumed origins of cortical evolution (Fig. 1 & S5). Several stable gradients represent highly specialized sensory modalities in the mouse cortex (Fig. 3). Other spatial features indicate a global functional progression from primary sensory areas towards more transmodal regions (Fig. 2).

### Intrinsic functional organization reflects the dual origin of the mouse cortex

We found that the principal gradient of functional connectivity in the mouse cortex reflects the spatial organization predicted by the dual origin theory (Fig. 1). This finding resonates with a recent study demonstrating a fundamental division of the mouse cortex into an archicortical and a paleocortical zone based on tract-tracing data (Goulas et al., 2019b). These authors also showed, that the dual organization was reflected in physical distance and gene expression data. Here, we used rsfMRI data acquired in vivo, and did not apply a cortical parcellation. This approach enabled us to complement the aforementioned findings with a gradient of functional connectivity, that captures the continuous progression from the two cortical origins, in addition to dividing the cortex into discrete zones. The principal gradient was reproduced across four data sets (Fig. S5) and provides robust evidence that the intrinsic functional organization of the mouse cortex in vivo is strongly influenced by the course of neocortical evolution from two primordial origins. Interestingly, studies on the intrinsic functional organization of the human cortex so far have not found a gradient clearly representing the dual origin axis. One possible reason is that it is harder to operationalize such a gradient based on simple geodesic distance from allo- and paleocortex, given the complex shape and folding of the human cortex. Another explanation could be that the human cortex is evolutionary further removed from its allocortical origins. It has been proposed that the massive expansion of transmodal areas in humans has *untethered* these areas from the influence of molecular gradients that constrain the organization of sensory regions (Buckner and Krienen, 2013). According to this hypothesis, the large transmodal regions of the human cortex are governed by different mechanisms and constraints than the sensory areas dominating the rodent cortex. Thus, while the distance from archicortex and paleocortex appears to be a constitutional feature of the rodent cortex, it might have been superseded by other organizational aspects in the human cortex.

### Sensory specialization is more pronounced than global processing hierarchies

Functional processing hierarchies are an established concept for the sensory cortex (Felleman and Van Essen, 1991) and have long been discussed as a more global idea, extending to transmodal regions (Mesulam, 1998). Recent studies on human cortical organization have consistently found a global gradient that ranges from sensory and motor cortex through heteromodal to transmodal association areas. This gradient describes a functional hierarchy of increasing abstraction in the temporal (Baldassano et al., 2017; Hasson et al., 2008; Lerner et al., 2011), semantic (Huth et al., 2016) and spatial domain (Peer et al., 2019), and in a meta-analysis of a multitude of task concepts (Margulies et al., 2016). The concept of global functional processing hierarchy has found less attention in the context of the mouse cortex. However, Fulcher et al (2019) recently described a sensory-to-transmodal progression in the average T1w:T2w values of cortical areas. T1w:T2w is commonly used as an in vivo proxy for microstructural differentiation, as it is thought to reflect intracortical myelin content (Glasser et al., 2014). Since the global functional hierarchy in humans has been shown to align with microstructural differentiation (Burt et al., 2018; Huntenburg et al., 2017), the T1w:T2w gradient described by Fulcher et al (2019) might point to a similar functional hierarchy in the mouse cortex. We therefore hypothesized that a sensory-to-transmodal pattern would be among the dominant gradients in the intrinsic functional organization of the mouse cortex. Indeed, we found aspects of a sensory-to-transmodal pattern represented in two separate gradients, Gradient 2 and 5 (Fig. 2A). We used the distance from primary sensory areas to operationalize the sensory-to-transmodal progression and found both gradients to be correlated to this measure (Fig. 2B & C). However, this organization differs in important ways from the respective findings in humans. The gradients representing the sensory-to-transmodal progression in the mouse cortex capture relatively less of the variance, and are more variable in their exact layout, than is the case for the human cortex. Instead of a single robust sensory-to-transmodal gradient, the variance associated with this type of organization is spread across at least two gradients – one emphasizing medial and one lateral transmodal regions, one clustering motor areas with the sensory, the other with the transmodal extreme. These gradients are also the least robust across data sets (Fig. S6A-C).

On the other hand, we found several, more robust gradients in the mouse cortex, associated with the specialization of different sensory modalities: Gradient 3 separates audiovisual from somatomotor areas, Gradient 4 captures the high level of specialization within primary somatosensory cortex (barrel field and nose vs. lower limbs and trunk), and Gradient 6 highlights primary auditory cortex (Fig. 3 & S6). This emphasis on sensory specialization in the mouse cortex is coherent with the importance of direct environmental input in this species. In the mouse, extensive parts of the cortex are dedicated to sensory processing and their most specialized sensory systems are most prominently represented. For example, mice heavily rely on their whiskers for spatial exploration and, correspondingly, the relevant barrel field area represents 19 % of their primary somatosensory cortex. Our findings suggest that the intrinsic *functional* organization of the mouse cortex, too, is heavily shaped by sensory processing, driven by thalamic inputs. As mentioned in the previous section, the dominance of sensory areas in the mouse is in contrast to the cortex of primates, especially humans, where hetero- and transmodal regions have massively expanded (Buckner and Krienen, 2013; Diamond and Hall, 1969). Humans have evolved to strongly rely on cognitive capacities that require abstraction from direct sensory input and are typically associated with computations in transmodal regions. It is plausible that such processing requires a strong emphasis on a sensory-to-transmodal organization, defined by cortico-cortical connections. In addition, human rsfMRI is typically acquired in an awake state, while our data has been acquired in lightly anesthetized rodents. It is possible that the anesthesia itself diminishes the dominance of the sensory-to-transmodal gradient, and that a similar result would be observed in anesthetized humans.

In summary, the intrinsic functional organization of the mouse cortex under anesthesia is heavily influenced by the specialization of sensory areas for processing of direct environmental input, while the awake human cortex is more strongly governed by a sensory-to-transmodal axis of information integration and abstraction. This sensory-to-transmodal axis is present, yet less pronounced in the functional organization of the mouse cortex.

### No evidence for an association between microstructural differentiation and functional connectivity gradients in the mouse cortex

Measures of microstructural differentiation have been shown to align with sensory-to-transmodal gradients in the human (Burt et al., 2018; Huntenburg et al., 2017), macaque (Burt et al., 2018) and mouse cortex (Fulcher et al., 2019). We did not find convincing evidence for such a relationship in our data. A modest correlation between the sensory-to-transmodal Gradient 2 and cytoarchitectonic type did not survive correction for multiple comparison (Fig. S4A & B). T1w:T2w as a proxy for intracortical myelin showed a weak correlation with the somatomotor-to-audiovisual Gradient 3, instead (Fig. S4C). One possible explanation for the overall weak correlation of functional gradients and microstructural measures that the mouse cortex is comparatively homogeneous with regard to microstructural differentiation. For example, the degree of “externopyramidization” – the thickness and architecture of supragranular vs. infragranular layers – increases progressively along the phylogenetic tree from rodents to marmosets, to macaque monkeys, to humans (Goulas et al., 2019a). Externopyramidization has been suggested to underlie the sensory-to-transmodal functional hierarchy (Goulas et al., 2019a, 2018; Sanides, 1962), and indeed, the dominance of this hierarchy appears to increase along the same phylogenetic trend: While we found no singular clear-cut sensory-to-transmodal gradient in the mouse cortex, such patterns have been described in secondary gradients of the marmoset (Buckner and Margulies, 2019) and macaque monkey (Margulies et al., 2016) based on connectivity data from tract-tracing experiments, and in the dominant principal gradient in humans (Margulies et al., 2016).

Yet, T1w:T2w has been associated with a sensory-to-transmodal hierarchy in a recent study of the mouse cortex (Fulcher et al., 2019), which made the lack of such an association in our data surprising. As the underlying T1w:T2w data is the same in both studies, much of the discrepancy might be explained by differences in methods. We compared functional connectivity gradients and T1w:T2w in a voxel-wise manner across the entire isocortex. Fulcher et al. (2019), by contrast, averaged T1w:T2w within cortical areas and compared those average values to other area-wise metrics. Averaging increases the signal-to-noise ratio of the T1w:T2w map, but it also obscures any variation within areas, potentially distorting the spatial distribution of T1w:T2w values. Both aspects might have contributed to differences in the outcomes. It is also important to keep in mind that the description of a gradient as “sensory-to-transmodal” is rather broad, so that the difference in correlation to T1w:T2w might partly derive from discrepancies between the sensory-to-transmodal gradient described by Fulcher et al. (2019), and our Gradient 2 and 5. In the context of the current study, we conclude that no substantial correlation was observed between two measures of microstructural differentiation, and the functional connectivity gradients analyzed.

### Strong links between dominant patterns of gene expression and functional connectivity gradients

A different approach to exploring the physiological underpinnings of functional connectivity gradients is through patterns of gene expression. We showed, that the somatomotor-to-audiovisual Gradient 3 and the sensory-to-transmodal Gradient 5 are highly correlated to dominant patterns in the expression of approximately 4000 genes in the mouse cortex (Fig. 4). Correlations between a principal component of gene expression and a sensory-to-transmodal gradient have previously been described in the human (Burt et al., 2018) and mouse cortex (Fulcher et al., 2019), in the same studies that reported associations of this gradient with T1w:T2w. The fact that we can reproduce the former, but not the latter result in our study, suggests gene expression to be a more robust counterpart to sensory-to-transmodal organization than T1w:T2w.

The strong correlation of somatomotor-to-audiovisual Gradient 3 and Gene PC3 suggests that the prominence of sensory specialization in the mouse cortex, discussed in previous sections, finds expression in a dominant molecular axis. When focusing on the genes most strongly associated with Gene PC3, we found an enrichment of ontologies related to cell cycle, cell differentiation and development. A possible interpretation is that the spatially distinct regulation of cell differentiation underlies the specialization of sensory processing in different areas of the mouse cortex. Taken together, our results reveal a strong link between the intrinsic functional organization, and the spatial variation in gene expression in the mouse cortex.

### A new perspective on the functional organization in animal studies

What is the advantage of the gradient approach for studying cortical organization? We have shown here that gradients can reveal organizational patterns that do not respect areal boundaries (most strikingly the principal gradient) and might be missed when imposing classic parcellation schemes. Without doubt, parcellations and the assignment of specific functions to discrete areas can be an extremely useful tool. And after all, we base much of the interpretation of the different gradient patterns on the known function of different areas. However, the functional assignment approach can also encourage oversimplified one-to-one mappings. This is particularly true in animal research, where highly controlled behavioral tasks and specialized measurement techniques sometimes prompt a mapping of individual task variables to small populations of cells. Gradient-based analyses, instead, produce superimposed representations that capture dominant patterns while also preserving information about other, coexisting functional dimensions. We apply this approach to resting-state functional connectivity data here, but believe that the general framework can be useful for many types of contexts and data. An excellent example is a recent study recording single cell responses in the posterior cortex of mice performing a visuomotor navigation task (Minderer et al., 2019). Using two-photon imaging, Minderer et al. show that task-relevant features are encoded in wide-spread gradients, rather than confined areas. Differences across the cortex are expressed in the weights of feature combinations and show, for example, that multimodal integration is most pronounced in regions where gradients encoding visual and motor aspects of the task overlap. The authors emphasize that we might miss much of the information in our data when confining our analyses to overly simplistic functional mapping, in particular when studying complex tasks or parts of the cortex that we know little about. We agree with this perspective and believe that gradient-based analyses are one important way of extracting more information from the increasingly complex data being acquired.

### Limitations and open questions

We used rsfMRI data to study the intrinsic functional organization of the mouse cortex in vivo. In contrast to post-mortem studies, this approach allows us to obtain data from the entire,intact and functioning cortex of each animal. However, imaging living animals comes with its own challenges. Physiological processes can introduce artifacts such as magnetic field distortions caused by air filled sinuses or movements associated with heart beat and breathing, which can only partly be removed during preprocessing. Most importantly, high-quality fMRI in rodents requires the animals to be lightly anesthetized. Here we used an optimized anesthesia strategy, aided by mechanical ventilation (Grandjean et al., 2014).While this protocol provides stable anesthesia and minimizes effects on spontaneous cortical activity, the anesthetic agents inevitably affect the physiological and brain state of the mice. This state is substantially different from the awake resting condition typical in human rsfMRI studies, and may underlie some of the observed differences between functional connectivity gradients in the two species. In particular, it is conceivable that the weaker expression of the sensory-to-transmodal gradient in the mouse cortex can partly be attributed to the effects of the anesthesia. It will be an interesting challenge for future research to understand the role of anesthesia in this context by employing rsfMRI data from awake mice and/or anesthetized humans. Another potential caveat is that the different types of data considered in our analyses are derived from cohorts of mice with different age and gender structures. For example, the Allen Mouse Brain Atlas is based on male mice only while our Main data set consists of only females. (However, the three test data sets consist of mostly males). The sexual dimorphism in the brain organization of C57Bl6 mice might therefore have influenced our findings. In order to reduce the chance that our results reflect phenomena specific to a single data set, rather than meaningful organizational principles, we cross-referenced the rsfMRI findings with independent measures and previous studies, and reproduce them across four data sets. While these data sets were acquired in different contexts, by different individuals and partly on different machines, it should be pointed out that they all stem from the same research group and are acquired under the same anesthetic protocol. It will be interesting to repeat the analyses across more diverse data sets.

Another point that warrants reflection is the interpretation of the gradients presented here. Although our analyses are largely data-driven and exploratory, our interpretations are obviously influenced by our assumptions and the literature we consulted. Alternative interpretations might be equally plausible. For example, the association of Gradient 1 with distance from paleocortex (piriform cortex) could also reflect the role of piriform cortex as the primary cortex of the olfactory system, one of the dominant senses in the mouse. Which of several possible interpretations will turn out to be most convincing, can only be settled by accumulating more evidence in future studies.

## Conclusion

We studied the intrinsic functional organization of the mouse cortex through gradients of functional connectivity. Our work complements previous studies on cortical organization in the mouse by focusing on superimposed, continuous gradient patterns extracted from in vivo rsfMRI data. We interpreted our findings in the context of the dual origin theory and cortical gradients described in other species, and conclude that a shift in gradient patterns occurs along the phylogenetic tree. Robust and dominant patterns in the mouse cortex display a strong link to the evolutionary origin of the neocortex, and highlight sensory specialization. In humans the emphasis has shifted to a core sensorimotor-to-transmodal pattern underlying a global hierarchy of information integration.

### CRediT author statement

JMH - Conceptualization, Methodology, Software, Validation, Formal Analysis, Writing - Original Draft, Visualization, Funding acquisition

LYY - Investigation

FM - Investigation

JG - Methodology, Software, Validation, Investigation, Resources, Data Curation, Writing - Review & Editing, Supervision, Funding acquisition

## Abbreviations

MRI: magnetic resonance imaging
rsfMRI: resting-state functional MRI
GE: gradient-echo
EPI: echo-planar imaging
FOV: field of view
MD: matrix dimensions
TR: repetition time
TE: echo time
Allen Mouse CCF v3: Allen Mouse Common Coordinate Framework version 3
Allen SDK: Allen Software Development Kit
PC(A): principal component (analysis)
SA: spatial autocorrelation
GO: gene ontology
FDR: false discovery rate
T1/2w: T1/2-weighted

## Acknowledgment

This project has received funding from the European Union’s Horizon 2020 research and innovation programme under the Marie Skłodowska-Curie grant agreement No 867458 awarded to JMH. LYY, FM, and JG were supported by the Singapore BioImaging Consortium core funds and the Singapore BioImaging Consortium awards awarded to JG (2016) and FM (2017).

