## Supplementary Material for "Gradients of functional connectivity in the mouse cortex reflect neocortical evolution"

### Supplementary tables and figures

| | Raw Data | Preprocessed Data | | Reference |
|---|---|---|---|---|
| | | URL | Selection Criterion | |
| **Main** | https://openneuro.org/datasets/ds001653 | https://doi.org/10.34973/1he1-5c70 | dataset='aes2' mediso_ctl='yes' | This article |
| **AD2** | https://central.xnat.org/data/projects/fMRI_AD_mouse2 | https://doi.org/10.34973/1he1-5c70 | dataset='AD2' mediso_ctl='yes' | Grandjean et al., 2016b |
| **AD3** | https://openneuro.org/datasets/ds001890 | https://doi.org/10.34973/1he1-5c70 | dataset='AD3' mediso_ctl='yes' | Mandino et al., 2019 |
| **CSD1** | https://central.xnat.org/data/projects/CSD_MRI_MOUSE | https://doi.org/10.34973/1he1-5c70 | dataset='CSD1' mediso_ctl='yes' | Grandjean et al., 2016a |

**Tab. S1.** Resources for raw and preprocessed data sets used in this study.

| GO Term | Description | $P_{corr}$ | Enrichment |
|---|---|---|---|
| GO:0051726 | Regulation of cell cycle | 0.026 | 7.46 |
| GO:0051270 | Regulation of cellular component movement | 0.037 | 1.70 |
| GO:0045595 | Regulation of cell differentiation | 0.032 | 1.32 |
| GO:0050793 | Regulation of developmental process | 0.042 | 1.27 |
| GO:0065007 | Biological regulation | 0.020 | 1.14 |

**Tab. S2.** Results of the gene ontology enrichment analysis for Gene PC3. FDR was controlled at 5% using the Benjamini-Hochberg method. Only ontologies with Pcorr ≤ 0.05 are shown.

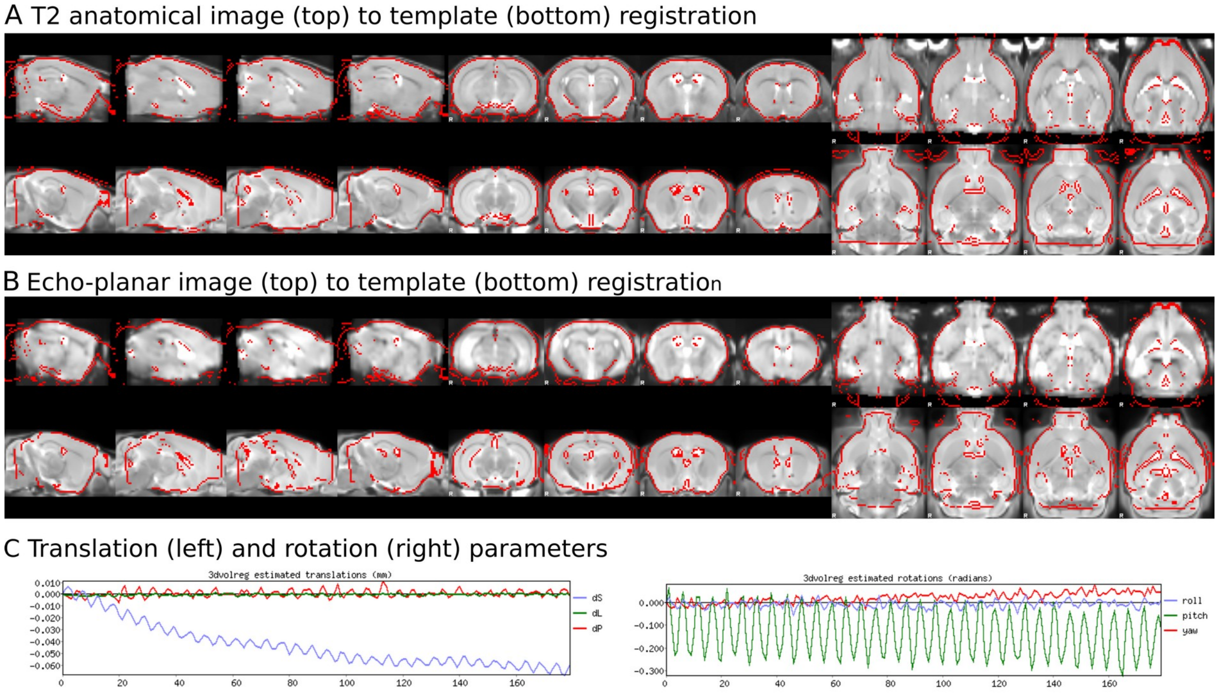


**Fig. S1.** Quality assurance procedure. A) Representative anatomical to template registration. B) Representative EPI to template registration. C) Representative motion traces (translation and rotations). Each run was visually inspected for motion and registration accuracy.

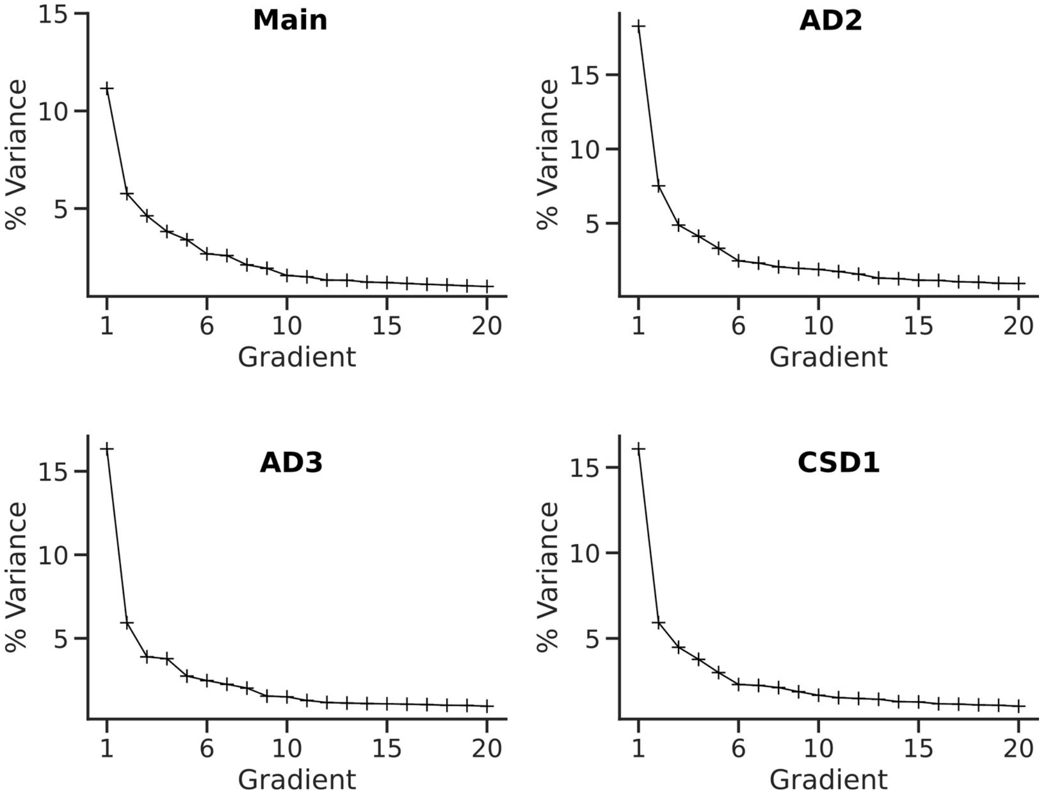


**Fig. S2.** Individual variance captured by the first 20 gradients in all four analyzed data sets. Note that the distributions are similar across data-sets. Data sets Main, AD2 and CSD1 show an elbow, i.e. a flattening of the variance curve, after Gradient 6. For AD3 a first elbow is visible at Gradient 4.

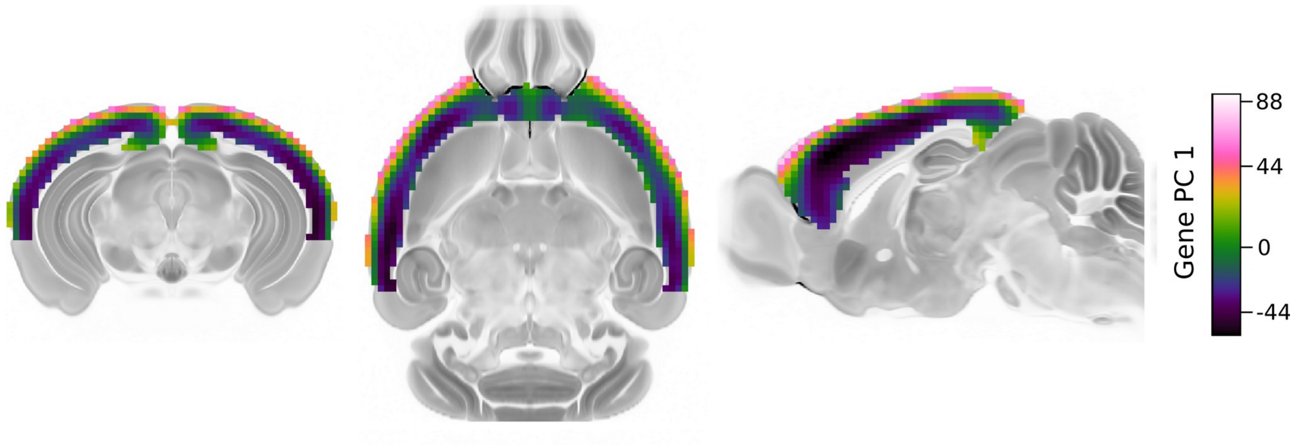


**Fig.S3.** Gene PC1 shows a cortical depth-dependent pattern.

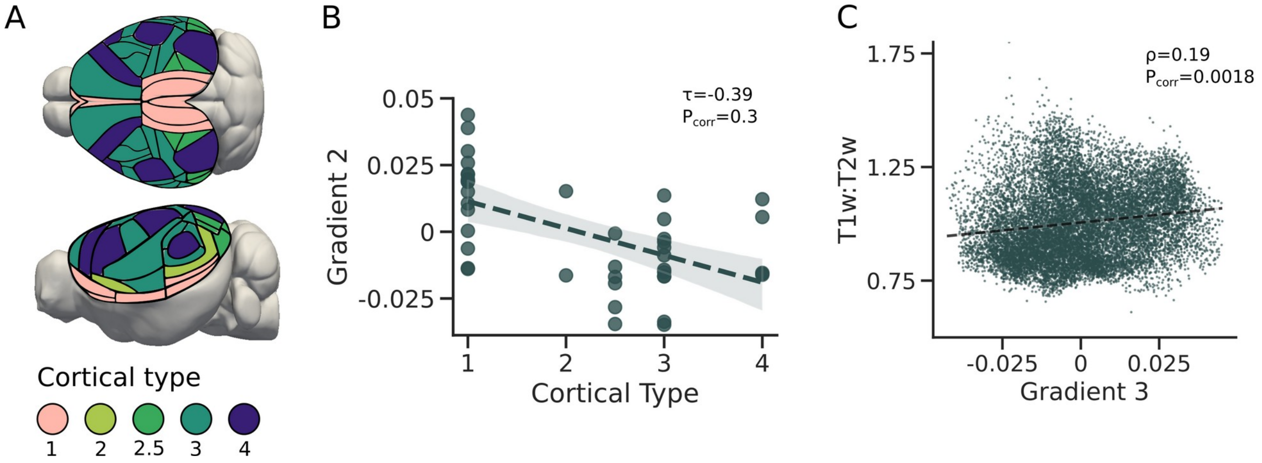


**Fig.S4.** Measures of microstructural differentiation. A) Nominal scale of cytoarchitectonic differentiation. B) Relationship between Gradient 2 and scale depicted in A (Kendall's τ=-0.39, $P_{corr}$=0.3). C) Relationship between T1w:T2w map and Gradient 3 (ρ=0.19, $P_{corr}$=0.0018).

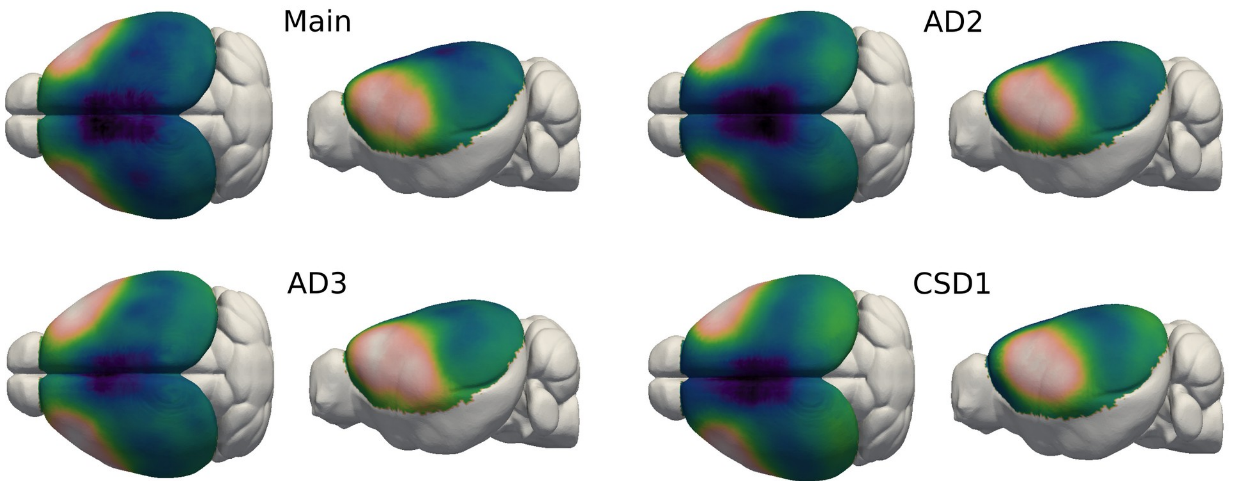


**Fig.S5.** The principal gradient of functional connectivity is qualitatively similar across all four data sets.

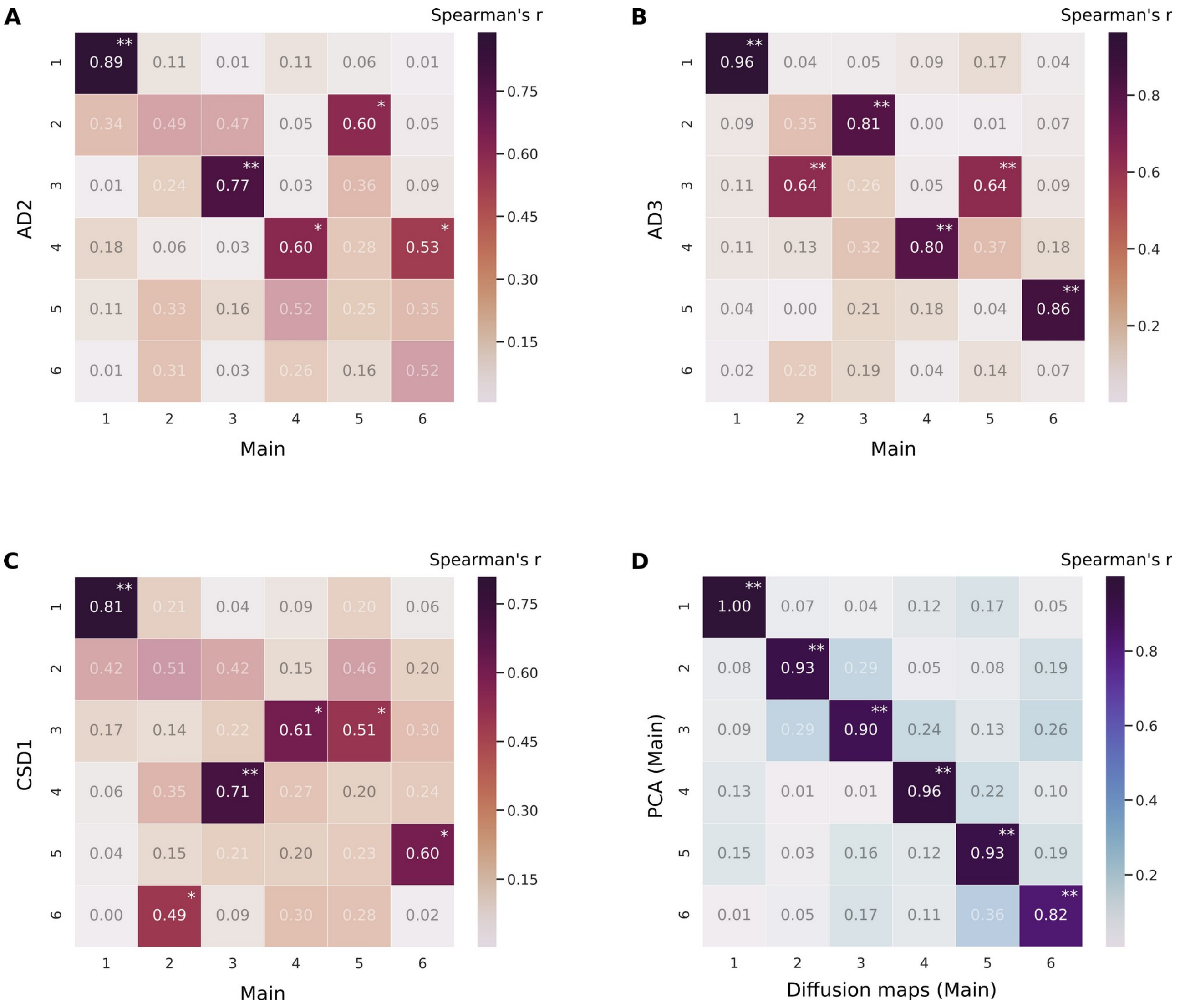


**Fig. S6.** Quantitative comparison between data sets and decomposition approaches. Values represent Spearman's rank correlation between Gradient 1-6 of the Main data set and Gradient 1-6 of data set AD2 (A), AD3 (B) and CSD1 (C), or components 1-6 derived via PCA of the Main data set (D). P-values were calculated through permutations tests and Bonferroni-corrected for multiple comparisons. * $P_{corr} \leq 0.05$, ** $P_{corr} < 0.0036$. Pairwise correlations with $P_{corr} > 0.05$ are faded out.
